# Prominent lateral spread of imaged evoked activity beyond cortical columns in barrel cortex provides foundation for coding whisker identity

**DOI:** 10.1101/137562

**Authors:** Nathan S. Jacobs, Ron D. Frostig

## Abstract

The posterior medial barrel subfield (PMBSF) of rat primary somatosensory cortex exquisitely demonstrates topography and columnar organization, defining features of sensory cortices in the mammalian brain. Optical imaging and neuronal recordings in rat PMBSF demonstrates how evoked cortical activity following single whisker stimulation also rapidly spreads laterally into surrounding cortices, disregarding columnar and modality boundaries. The current study quantifies the spatial prominence of such lateral activity spreads by demonstrating that functional connectivity between laterally spaced cortical locations is actually stronger than between vertically spaced cortical locations. Further, the total amount of evoked activity within and beyond single column boundaries was quantified based on intrinsic signal optical imaging, single units and local field potentials (LFPs) recordings, revealing that the vast majority of whisker evoked activity in PMBSF occurs beyond columnar boundaries. Finally, a simple two layer artificial neural network model of PMBSF demonstrates the capacity of extra-columnar evoked activity spread to provide a foundation for accurate whisker stimulus classification that is robust to random scaling of inputs and local noise. Indeed, classification performance improved when more of the lateral spread was included in the model, providing further evidence for the relevance of the lateral spread.

## Introduction

The rat primary somatosensory cortex, especially the posterior medial barrel subfield (PMBSF) representing the large, moveable whiskers (vibrissae), exquisitely demonstrates topography, a defining feature of sensory cortices in the mammalian brain. Each large whisker found on the rodent snout corresponds to a unique cytoarchitectural unit in layer IV of PMBSF known as whisker barrel. Topographic organization is further supported by columnar organization in which neurons above, below, and within a barrel in layer IV respond preferentially to the same whisker. Topographic and columnar organization of PMBSF is supported by vertically oriented excitatory connections within barrel columns^1^. PMBSF thus adheres to clear topographic and columnar principles of organization.

In addition to columnar and topographic boundaries, PMBSF also exhibits lateral spread of activity with prominent spatial footprints that span several mm in rats. Spatially broad profiles of activity evoked by “point” stimuli such as a single whisker are a ubiquitous and notable feature of sensory cortex^2-20^ (reviewed in [20]). In PMBSF, lateral spread of activity is supported by an underlying plexus of horizontal intracortical projection fibers^14,21-23^.

The functional contributions of lateral spread in PMBSF have only recently begun to be explored^20^. The lateral spread has been linked to integration of cortical responses following multi-whisker stimulation^16^ and support whisker representations that are independent of absolute response magnitude and invariant to large changes in stimulus amplitude^19^. Importantly, both studies found a primary role for whisker-evoked activity *beyond* cortical columns, rather than peak responses within columnar boundaries. These beyond-the-column findings were surprising as it is commonly assumed that only the strongest (or peak) response is the important variable for cortical processing and function. These findings, therefore, warrant a further study into the spread of cortical activity laterally beyond the column. The current study quantifies the prominence of the lateral spread beyond columnar boundaries using a functional connectivity analysis and cumulative response calculations. In addition, a simple artificial neural network model was used to demonstrate how activity patterns produced by lateral spread can support robust whisker identity coding. These findings provide further evidence for the importance of lateral spread beyond peak activity within PMBSF.

## 2 Materials and Methods

### Subjects

All in vivo procedures were in compliance with the National Institutes of Health guidelines and reviewed and approved by the University of California Irvine Animal Care and Use Committee. Subjects were adult male Sprague–Dawley rats. Rats were inducted with a bolus intraperitoneal injection of sodium pentobarbital (55 mg/kg b.w.) and maintained with supplemental injections as needed throughout the day. For intrinsic signal optical imaging (ISOI) data 37 rats were used. For multi-depth electrophysiology recordings a different set of 7 rats were used.

### Whisker stimulation

Whisker stimulation was restricted to only the right snout. Single whisker stimulation targeted the C2 whisker located centrally on the whisker pad **(Fig. 3A)**. As in previously established protocols, the stimulation of only whisker C2 was achieved with a copper wire probe attached to a computer-controlled stepping motor. Five deflections were delivered at 5 Hz rate for total time span of 1 s. Each deflection displaced whisker C2 approximately 1 mm along the rostro-caudal direction at an approximate speed of 0.25 mm/ms (whisker probe position stabilized within 10 ms of initiation of movement) at a distance of approximately 5 mm from the skin.

### Intrinsic signal optical imaging

Intrinsic signal optical imaging (ISOI) was used for high-spatial resolution, wide field-of-view mapping of the total cortical activation spread evoked by whisker stimulation. Imaging was conducted with a 16-bit CCD camera (Cascade 512B II; Photometrics, Tucson, AZ) combined with an inverted 50 mm lens plus extenders. The camera’s field-of-view was a 7.42×7.42 mm cortical region, mapped onto a 256×256 pixel array. For future alignment of data files collected within the same rats as well as across rats, the field-of-view neuroaxis was oriented the same in every rat, plus the field-of-view remained constant across data files within each rat. The CCD camera was focused 600 μm below the cortical surface before the start of data collection to minimize contributions from surface blood vessels and maximize contributions integrated across the upper cortical layers. The imaged cortical region was continuously illuminated with a red LED (635 nm max, 15 nm full width at half-height). Imaging frames were captured at 10 Hz rate (i.e., 1 frame = 100 ms exposure time), and each imaging trial lasted 15 s. Onset of whisker stimulation occurred 1.5 s into the trial. A block of 64 trials was collected per whisker stimulation condition, with an intertrial interval averaging 6 s and ranging randomly between 1–11 s and thus an average of 21 s between the onset of consecutive stimulus deliveries. The 64 trials in a block were then summed and the summed data collapsed into 500-ms frames (referred to hereafter as a data file) to increase the signal-to-noise. For each data file, activity for each 500-ms post-stimulus frame was converted to fractional change relative to the 500-ms frame collected immediately prior to stimulus onset on a pixel-by-pixel basis. The current data set focused on “initial dip” frames in which the first local minimum of the ISOI signal occurs. C2 whisker barrel column and PMBSF boundaries were estimated using a representative map of whisker barrels derived from cytochrome-oxidase stained sections.

### Electrophysiology

Multi-site, extracellular recordings were acquired using 32-channel arrays with an 8×4 design consisting of 8 recording locations each of which had four depths targeting layers 1, 2/3, 4, and 5 (**Fig. 2A).** Microelectrode arrays were made from insulated 35 um tungsten wire (California Fine Wire, Grover Beach, CA). that were blunt cut and threaded in groups of four through polyimide guide tubes spaced 0.5 mm apart. Mean impedance of electrodes was 153 kΩ ± 55 (measured with IMP-2, Bak electronics, Sanford, FL). Raw signals starting 1 s before and ending 1 s after stimulus onset (total of 3 s per trial) were amplified and digitized at a 22 kHz sample rate (SnR system, Alpha Omega, Nazareth, Israel). Distance to C2 whisker barrel was measured in cytochrome oxidase stained tissue sections. For PSTHs, 1 ms time bins were used.

Post-processing was done using custom MATLAB scripts. Raw traces were band-pass filtered for local field potentials (LFP, 1–300 Hz) or spikes (300–3k Hz) using a two-pole Butterworth function. Trials with electrical noise (5.32% of trials) were excluded from trial averages. For the few bad channels in arrays (5.36% of channels overall, equivalent to 1.6 channels per array), trial averages from adjacent channels at the same cortical depth were averaged. In trial averaged LFP, mean baseline values 50 to 0 ms before stimulus onset were subtracted. A Gaussian filter was used to remove electrical noise near 60 Hz. Data was down-sampled to a 10 kHz sample rate for further analysis.

For z-normalized LFP data in **Fig. 1**, responses were divided by the standard deviation of signal in the 10 ms time period before whisker stimulation on a trial-by-trial basis. This resulted in values that indicated the number of standard deviations from baseline noise a given response was.

**Figure 1.**
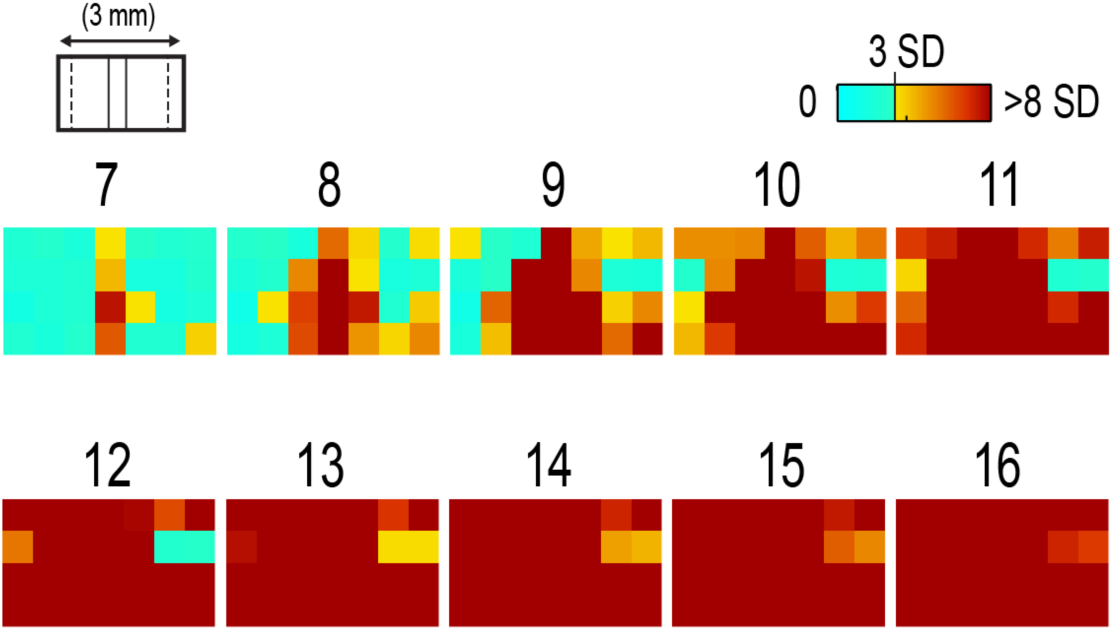
Beyond columnar activity. Time series of single whisker (C2) evoked LFP plotted as z-scores of baseline signal (mean, n=6, data set from [19], see methods for details). Multi-site recordings were taken from a planar section of rat PMBSF penetrating through the depth of cortex. Each frame consists of 7 ×4 recording locations in a multi-electrode recording array sampling from cortical layer 1 (top of image), 2/3, 4, and upper layer 5 (bottom of image). Laterally, the array extends from just beyond PMBSF rostrally (left edge of image) to just beyond PMBSF caudally (right edge of image). Initial evoked activity is seen across vertical column of tissue consistent with traditional columnar boundaries. This brief (1 ms) initial pattern is rapidly transformed to extensive lateral spread of cortical responses into surround cortex. Within milliseconds, evoked activity > 3 SD above pre-stimulus noise is seen across all sampling locations. *Inset, upper left, simplified schematic of electrode array positions (for detail see Fig. 1A). Dark lines denote columnar boundaries of C2 whisker barrel, dotted lines indicate PMBSF boundaries.*

### Whisker identity coding model based on spatial pattern of activity

Whisker coding based on mesoscopic patterns of evoked activity was assessed using a two layer artificial neural network consisting of a 5x5 input layer (“PMBSF” with 25 “whisker barrels”) and a single output unit (downstream “reader” neuron).

The network architecture was similar to a perceptron^24^. Input units were only connected to the output unit. There was no feedback onto the input layer. Synaptic weights were randomly set to a value between −1 and 1. The degree and arrangement of connectivity between the input and output layer was varied to produce four network types (see **Fig. 4C**): 1) single connection between input layer and output unit, 2) four adjacent units of input layer connected to output unit, 3) four maximally spaced units of input layer connected to output unit, and 4) all units of input layer connected to output unit. This set of network architectures was chosen because it included variations in the degree of connectivity (1 vs 4 vs 25 inputs) as well as variations in the heterogeneity of connections (adjacent vs spaced inputs).

Each trial consisted of two time steps: 1) activation of input layer and 2) activation of output unit. Input activity was set by empirically derived activity patterns (see below) with values ranging between 0 and 1. Output unit activity was binary (0 or 1) and calculated using a thresholded, linear activation function with no bias constant and threshold = 0.

Input activity patterns were empirically derived from previously published ISOI data^16^. A response decay rate function was derived from average whisker C2-evoked ISOI “initial dip” data obtained by averaging results from 37 rats (**Fig. 4A**, left panel). Four linear sections starting at the peak response and extending radially outward each separated by 90 degrees were averaged, peak-normalized, and fitted with a two component exponential function (**Fig. 4A**, middle panel): 0.2*exp(-0.07*(x*s)) + 0.83*exp(-0.014*(x*s)), where s=21 to convert from spatial scale of empirical data (1 pixel = 28 um) to spatial scale of model input layer (1 pixel = 0.5 mm, approximate distance between barrels in rat PMBSF). The decay rate function was used to produce 25 different activity patterns with peak activation over each of the 25 pixels of the 5×5 input layer (**Fig. 4A**, right panel). Lastly, in order to model changes in stimulus intensity, on every trial the activity pattern was also uniformly scaled by a random factor between 0 and 1 pulled from a uniform distribution function (rand() function in MATLAB). Uniform scaling of mesoscopic response patterns in PMBSF across a wide range of stimulus intensity has previously been shown^19^.

Each trial was randomly selected to either include presentation of the target input (C3, with peak activation over the “barrel” at the very center of the 5x5 input layer) or an off-target input (all other whisker inputs). For trials of off-target inputs, one of the 24 non-target input activity patterns was randomly selected. Thus, half of all trials included the target input and chance performance was 50% accuracy. Performance (% of trials accurately classified as on or off target) was assessed in blocks of 100 trials called a “session.” Training consisted of 20 sessions (total of 2,000 trials) and testing consisted of 10 sessions (total of 1,000 trials). Synaptic weights were updated during training (trials 1-2,000) using the delta rule: _ij_ = * (t_j_ – y_j_) * x_i_, where is synaptic weight, *i* is the presynaptic unit, *j* is the postsynaptic unit, is the learning rate constant, *t* is the target response, *y* is the actual response, and *x* is unit activity. The simulation was repeated for 100 networks (per network type).

Network simulations were repeated for three conditions: 1) no noise, 2) noise, and 3) noise with complex classification task. The no noise condition is described above. For the noise condition, independent noise (random value between −0.1 and 0.1) was injected into each unit of the input layer on each trial. For comparison, peak values in input patterns ranged between 0 and 1 with a theoretical mean of 0.5. Thus, the maximum signal to noise ratio for peak values under these conditions was 10, with a theoretical average of 5 and even lower signal to noise ratios for all off-peak values.

For the last condition, noise was again added and the target input was made more challenging – the output unit had to respond to both the C3 and A5 inputs (while still correctly rejecting all other inputs).

MATLAB source code at: https://zenodo.org/badge/latestdoi/87992624.

### Statistics

Cross correlations, modeling, and descriptive statistics were performed using custom MATLAB scripts. All parametric statistics were performed in SYSTAT version 11.

For cumulative response magnitudes, responses between 0 mm and 0.22 mm, 0.5 mm, 1.0 mm, or 1.5 mm from location of peak activity were integrated. For LFP and MU data, response value at 0.22 mm was interpolated (linear interpolation) from response values at 0 mm and 0.5 mm and the trapezoid method of integration used for cumulative response calculations.

Cross-correlations of evoked LFP with +/- 50 ms lag times were run between pairs of recording locations of an 8x4 array of electrodes using MATLAB’s xcorr() function. Each recording location was used as a seed and cross correlations run with locations within the same vertical column (for vertical connectivity measure) or within the same horizontal row (for horizontal connectivity measure).

For box and whisker plots in **Fig. 2**, solid line indicates the median, outer bounds of box indicate upper and lower quartiles, whiskers show minimum and maximum values (excluding outliers), and dots represent outliers using the following default formula for box plots in MATLAB: Maximum whisker length w is set to 1.5 times the interquartile range. Outliers are any data points greater than q3 + w(q3 - q1) or smaller than q1 - w(q3 - q1), where q1 and q3 are the 25th and 75th percentiles, respectively.

**Figure 2.**
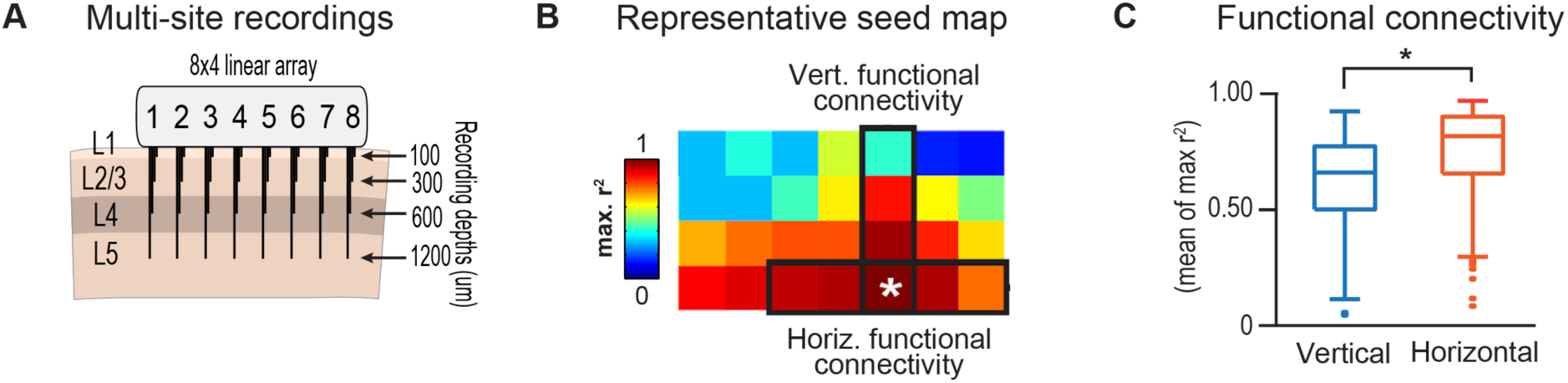
Horizontal and vertical functional connectivity in barrel cortex. **(A)** Multi-electrode array design to sample from most cortical layers (up to 1.2 mm deep) and across the full breadth of rat barrel cortex (3.5 mm across). **(B)** Cross correlations on C2 evoked LFP data from [19] were run for each recording location. Mean r^2^ values within nearby horizontal recording locations were interpreted as “horizontal” functional connectivity. Mean r^2^ values for three other electrodes within the bundle were interpreted as “vertical” functional connectivity. **(C)** Group statistics show stronger functional connectivity along the horizontal axis (within layer) as compared to vertical ais (within column) for single whisker stimulation (C).

## 3 Results

### 3.1 Lateral spread of whisker evoked cortical activity

The stimulation of a single whisker in rats evokes a prominent lateral spread of activity through PMBSF and into surrounding cortices. An example of single whisker evoked LFP in PMBSF is shown in **Fig. 1**, which shows response magnitudes in standard deviations from baseline noise (see methods for details). Every sampling location - which included electrodes from 4 different cortical layers extending beyond the boundaries of PMBSF - recorded evoked LFP that was over 3 standard deviations above baseline noise. These data suggest that evoked activity is only constrained to a cortical column for 1 ms following single whisker stimulation, before spreading laterally beyond columnar boundaries and engaging a broad volume of cortex.

Robust horizontal functional connectivity would be consistent with the prominent lateral spread of evoked activity, however this has not been explicitly tested before. Therefore, functional connectivity along vertical and horizontal domains in PMBSF was compared using cross-correlations of multi-site recordings of single whisker (C2) evoked LFP (**Fig. 2**). Correlation coefficients for cross-correlations with a seed location were binned vertically or horizontally (see representative seed map in **Fig. 2B**) and then averaged for all seed locations. Mean peak r^2^ values were significantly *higher* for horizontal connectivity compared to vertical connectivity (*p=0.001 paired t-test*; **Fig. 2C**). These data suggest a robust horizontal connectivity that is stronger than the well-established vertical connectivity in PMBSF.

### 3.2 Prominent cumulative evoked activity beyond columnar boundaries

Stimulation of a single whisker in rats evokes responses within columns of cortical tissue in PMBSF that rapidly spreads laterally into surrounding cortices (**Fig. 1**). The cortical response to single whisker stimulation can thus be broken into two components-peak responses localized within cortical columns (strong response, small volume, **Fig. 3**, blue responses) and a broad profile of weaker responses (weak response, large volume, **Fig 3**, orange responses). To determine whether most single whisker evoked activity in PMBSF occurs near peak responses (i.e., within columnar boundaries) or in surrounding cortices (i.e., beyond columnar boundaries), cumulative response magnitudes were calculated.

**Figure 3.**
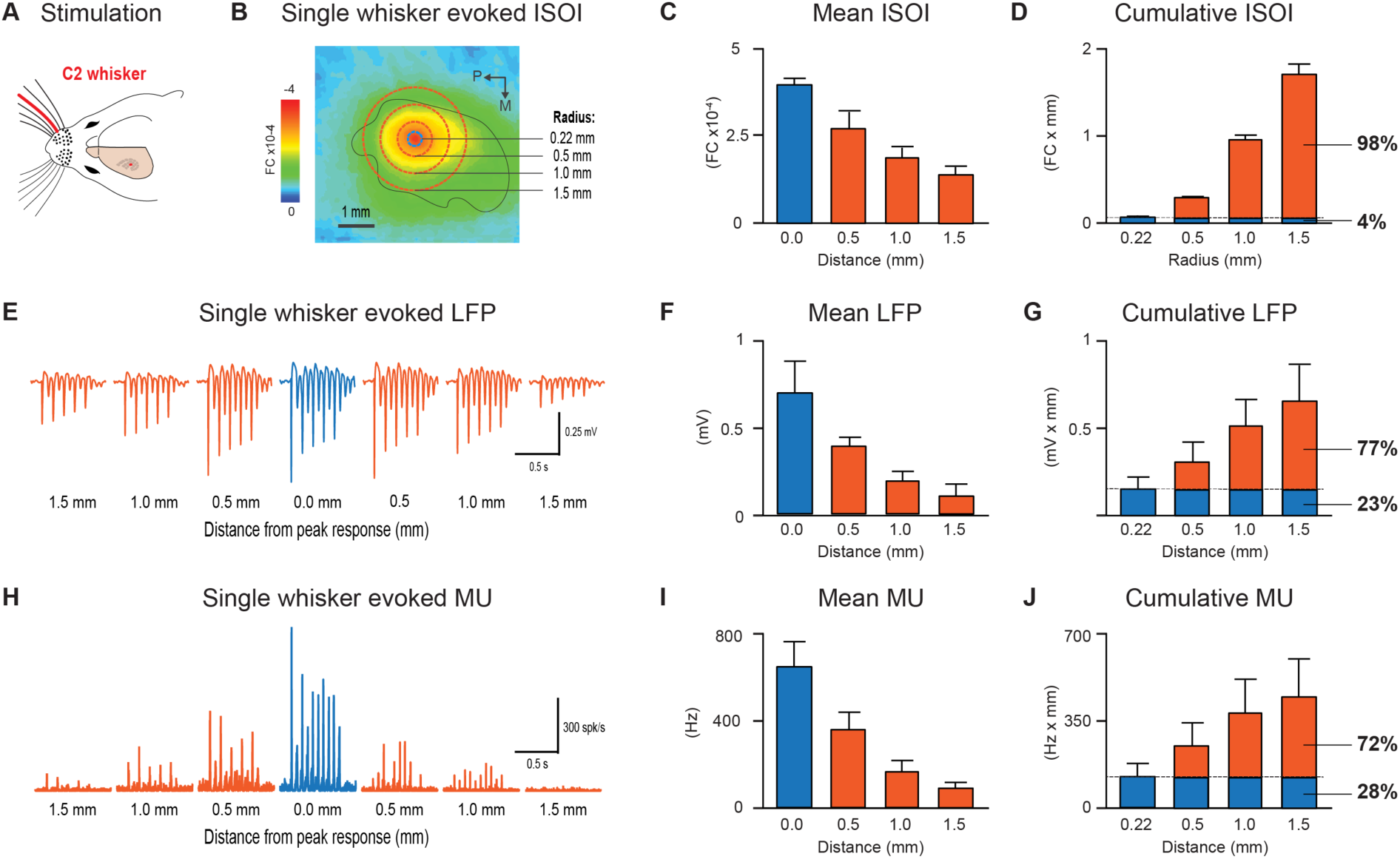
Single whisker evokes more activity *outside* barrel column. **(A)** Schematic of single whisker and anatomical representation in PMBSF. **(B, E, H)** Single whisker evoked activity in PMBSF measured by ISOI “ initial dip” signal (B, data set from [16]), LFP (E, data set from [19]), and multi-unit activity (H, data set from [19]). *FC*, *fractional change.* **(C, F, I)** Mean response magnitudes at 0, 0.5, 1, and 1.5 mm from location of peak c2 evoked response for ISOI (C), LFP (F), and MU activity (I). **(D, G, J)** Cumulative responses at 0.22, 0.5, 1.0, and 1.5 mm from location of peak C2 evoked response for ISOI (D), LFP (G), and MU activity (J). For two-dimensional ISOI data (D), responses were integrated within specified radius. For one-dimensional electrophysiology data (G, J), responses were integrated between 0 mm and specified distance from peak response. Responses at 0.22 mm were estimated using linear interpolation. **(E, H)** Representative single whisker evoked LFP (E) and multi-unit PSTHs (H) at increasing distances from peak response. *Recording depth = 600 um. Solid black line in B denotes PMBSF boundary.*

Following single whisker C2 stimulation (**Fig. 3A**), cumulative response magnitudes (response magnitude integrated across space) were calculated for increasing distances/areas from the location of peak responses for intrinsic signal optical imaging (ISOI) initial dip data (**Fig 3D**, n=37, data set from [16]), local field potential (LFP, **Fig. 3G**, n=6, data set from [19]), and multi-unit activity (MU, **Fig. 3J**, n=6, data set from [19]). A distance/radius of 0.22 mm was used as the standard demarcation for columnar boundaries based on the 0.15 mm^2^ area of the C2 whisker barrel^9^. Cumulative response magnitudes were calculated up to a distance/radius of 1.5 mm, roughly corresponding to the boundaries of PMBSF as measured from the C2 whisker barrel. On average, the cumulative response between 0 and 0.22 mm from peak responses (ie total activity within columnar boundaries) was only 4%, 23%, and 28% of the cumulative response between 0 and 1.5 mm from peak responses (i.e., total activity within and beyond columnar boundaries, up to 1.5 mm) for ISOI, LFP, and multi-unit data sets, respectively. The lateral spread of evoked multi-unit activity tends to be more spatially restricted than evoked LFP^14^ and ISOI activity^16^ which could explain the range in cumulative response magnitudes beyond columnar boundaries. For both activity measures, these data show that the vast majority of whisker-evoked activity (as much as 98%, for ISOI data) occurs beyond columnar boundaries. These percentages would be even smaller if the entire lateral spread of evoked activity, which extends beyond PMBSF boundaries^14^, were included.

### 3.3 Robust whisker identity coding using evoked activity spread

What is the purpose of all this extra-columnar activity? A possible role in providing a robust foundation for whisker coding is explored.

Robust coding of whisker identity with spatial activity patterns was tested using a two layer artificial neural network model of PMBSF and a downstream “reader” neuron. The input layer of the network was a 5×5 grid representing a simplified model of PMBSF with 25 whisker barrels separated by 0.5 mm (**Fig. 4A**, right panel). The pattern of activity across the input layer was empirically derived from C2-evoked ISOI “initial dip” data^16^, which was used to create a standard curve for the decay rate of single whisker evoked point spreads in rat PMBSF (**Fig. 4A**, middle panel). The simplified 5x5 patterns of activity were used to demonstrate an inherent ambiguity in whisker evoked columnar activity and, combined with downstream “reader” output units, to test whether spatial patterns of activity could support a robust whisker identity coding mechanism.

**Figure 4.**
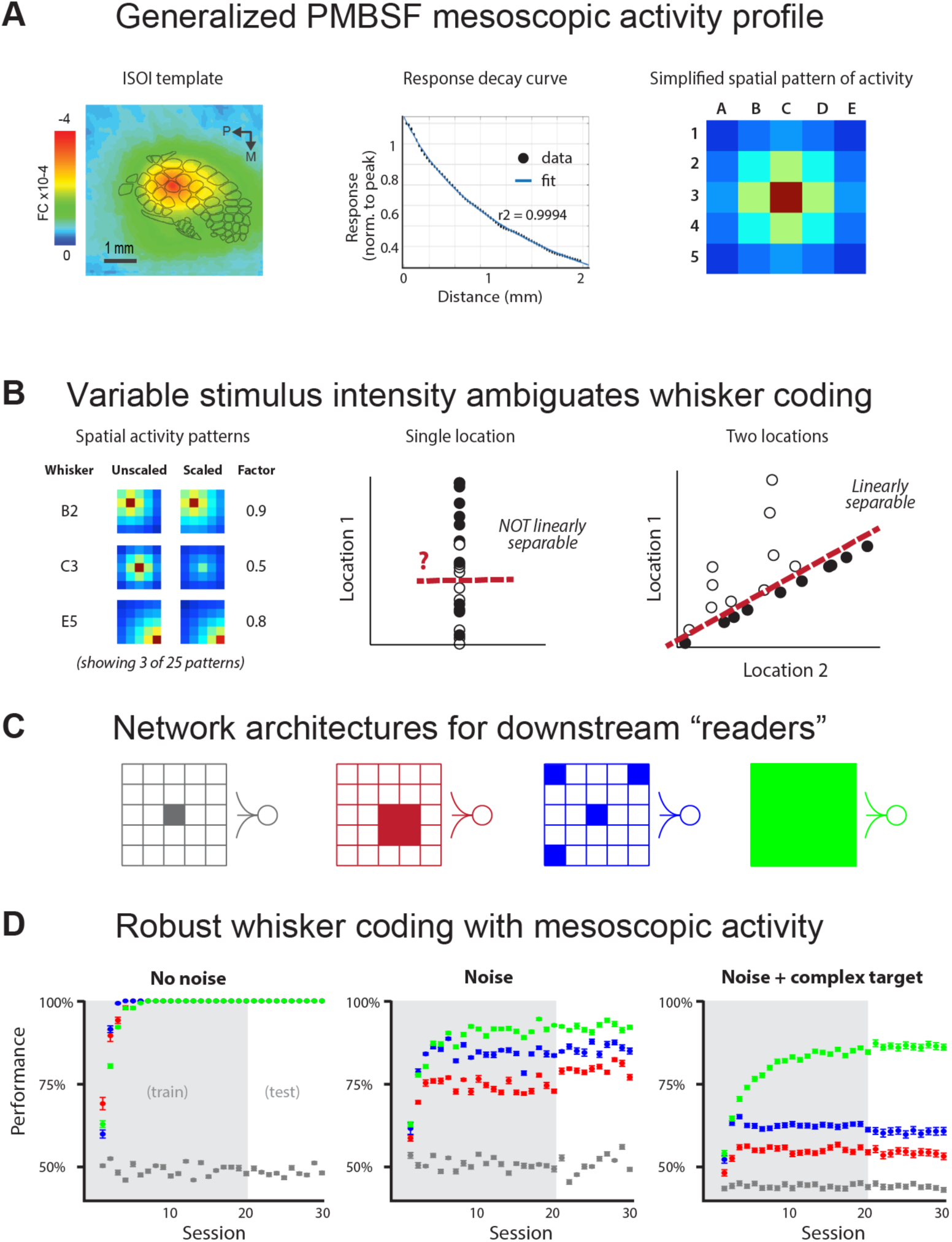
Spatial patterns of activity support robust whisker identity coding scheme. A simple two layer neural network model was used to test the robustness of whisker coding based on large spatial patterns of cortical activity. **(A)** Simplified cortical activity patterns were derived from mean whisker C2-evoked ISOI “initial dip” data from rat PMBSF (n = 37, data set from [16]). Left panel, mean ISOI “initial dip” data used to fit the decay rate function. Middle panel, ISOI response magnitudes plotted against cortical distance from peak response (black circles) with overlay of fitted response decay function (blue line). Right panel, example of a 5x5 stimulus pattern (1 pixel = 0.5 mm) generated by the response decay function. *Scale bar, left panel = 1mm.* **(B)** Before being used as inputs into the model, stimuli were multiplied by a randomized scalar to capture the variable intensity of whisker stimuli (left panel). Each whisker stimulus would therefore vary greatly in absolute response magnitude from trial to trial while the relative pattern of activity remained exactly the same, consistent with a previous report of invariant relative spatiotemporal profile of activity across large ranges of stimulus intensities^19^. Given the slow response decay rate, these conditions made it effectively impossible to classify whisker stimulus identity (e.g., whisker C3) using responses at any one location (middle panel). If responses from two locations are considered, the classification problem is greatly simplified (right panel). **(C)** Gray, reader unit connected to single location of input layer located at the peak response for the target stimulus (C3). Red, reader unit connected to four adjacent units of the input layer all within close proximity to the peak response for target stimulus. Blue, reader unit connected to four units of the input layer, one at the location of peak response for target stimulus and three additional units at maximal distances from the first. Green, reader unit connected to all units of the input layer. **(D)** Whisker classification performance for each reader type. Classification accuracy for all readers was tested in three conditions of increasing difficulty: no noise (left panel), noise (middle panel), and noise with complex target where readers had to respond to both the C3 whisker pattern as well as the A5 whisker pattern (right panel, ‘complex target’). *Error bars = SEM.*

If input patterns are scaled by a random factor across trials (**Fig. 4B**, left panel), there is no simple way of classifying whisker identity using input activity from just one location of the input layer (**Fig. 4B**, middle panel). This would be analogous to receiving afferents from neurons in a single cortical column. The difficulty in classifying whisker stimulus identity using activity from a single cortical location arises from the overlap in response magnitude distributions produced by a range of possible whisker stimulus intensities. For example, weak stimulation of the C2 whisker may produce a weaker response in the C2 barrel than strong stimulation of the C3 whisker. A simple way to resolve this inherent ambiguity is to integrate activity from multiple locations across PMBSF (**Fig. 4B**, right panel).

The output unit of the neural network modeled a downstream “reader” neuron decoding which whisker was stimulated. Four network architectures with varying degrees of connectivity between the input layer and the output unit were investigated (**Fig. 4C**). Classification performance of each network architecture was compared across three conditions of increasing difficulty-no noise (**Fig. 4D**, left panel), noise (**Fig. 4D**, bottom middle panel), and noise with a complex target (**Fig. 4D**, right panel). Results show that performance above chance (50%) required readers to be connected to multiple units of the input layer (**Fig. 4D**, compare gray circles to all other circles). Performance improved if the reader was connected to more heterogeneous units of the input layer (**Fig. 4D**, compare blue and red circles). Performance was best if the reader was fully connected to the entire input layer (**Fig. 4D**, compare green and blue circles). These results show that using the full spatial profile of activity (**Fig. 4D**, green network and circles) can support whisker coding that is robust to random scaling of input patterns, local noise designed to produce an average signal to noise ratio of 5 for peak responses and 2.5 for more distal responses (+1.5 mm from peak response, see methods for details), and supports learning a classification scheme in which the output unit responds to either of two non-adjacent whiskers but not to any others (with 85% accuracy for fully connected output unit, **Fig. 4D**, right panel, green circles).

In summary, these results suggest that stimulating a single whisker engages the entirety of PMBSF (**Fig. 1**) with stronger horizontal functional connectivity than vertical functional connectivity (**Fig. 2**), evokes more activity beyond columnar boundaries than within (**Fig. 3**), and produces spatial patterns of activity that support highly robust whisker identity coding (**Fig. 4**).

## 4 Discussion

Single whisker stimulation initially evoked activity within a cortical column^25^ that ignites an expansive profile of activity extending across and beyond PMBSF^14^. This finding was demonstrated again here (**Fig. 1**) and further supported by a horizontal functional connectivity that was stronger than vertical functional connectivity (**Fig. 2**). Importantly, the current findings demonstrate that rather than being the minority response, weaker, off-peak responses beyond columnar boundaries actually constitute the large majority of whisker evoked activity (**Fig 3**). This raises an important question - what is the functional role of the majority of whisker evoked activity in PMBSF?

Lateral spread of evoked activity has been linked to two important sensory functions in PMBSF, multi-whisker integration^16^ and invariance^19^. The current findings suggest a possible third function of prominent lateral spread of evoked activity - the robust coding of whisker identity. To test this, a simple two layer artificial neural network similar to the perceptron^24^ was used with spatially organized input patterns derived empirically from ISOI data (**Fig. 4A**). Spatial patterns of activity across PMBSF resolve an inherent ambiguity introduced by variable stimulus intensity (**Fig. 4B**). Increasing the degree and variety of sampling of input patterns improved the accuracy of whisker coding (**Fig. 4C-D**). The current findings reinforce the notion that coding is optimal with large, heterogeneous neural populations^28^ and demonstrate that even highly simplified spatial activity patterns can support highly robust whisker coding. The capacity for classifying whisker input sources should also hold for multi-whisker stimuli, provided they evoke unique spatial patterns of activity.

It should be noted that it is well known that even simple neural network architectures such as the perceptron can exhibit robustness to noise^26^ and perform classification tasks^27^. In the current study, these findings were simply replicated with a specific use case to demonstrate the ease with which whisker evoked activity patterns in PMBSF can be used to code for whisker stimulus identity.

The current study attempted to answer a simple question - where does most whisker-evoked activity occur within PMBSF? Finding that the large majority of whisker evoked activity occurs beyond columnar boundaries (spatial prominence) raises a second question – what is the role of this extra columnar spatial prominence? In addition to previously discovered roles in multi-whisker integration^16^ and invariance^19^, the current findings suggest that spatial patterns of activity may play a more general role as a foundation for robust whisker identity coding. Together, these findings demonstrate that spatially prominent, extra-columnar whisker evoked activity is a primary response feature in PMBSF.

If weaker off-peak lateral evoked responses are unimportant in PMBSF, why do they exist at all? And why should cortex maintain a system of horizontal projections to support them, rather than supporting only columnar activation? The current study answers this question in two important ways. First, we show with three complimentary neurophysiology data sets that despite weaker absolute response magnitudes, extra-columnar activity actually constitutes the large majority of whisker evoked activity in PMBSF. Second, we used a simple artificial neural network to show that spatial patterns of activity provide a highly robust foundation for whisker coding that is independent of absolute response magnitudes. Together the current results suggest that whisker evoked activity beyond columnar boundaries are as important for PMBSF as the columnar response and therefore the two should not be separated.

## Acknowledgements

Funding sources: This work was supported by the NIH-NINDS NS-066001 and NS-055832, and The Center for Hearing Research NIH Training Grant 1T32DC010775-01.

